# Characterization of mating type genes in heterothallic *Neonectria* species with emphasis on *N. coccinea, N. ditissima,* and *N. faginata*

**DOI:** 10.1101/2020.01.22.915686

**Authors:** Cameron M. Stauder, Jeff R. Garnas, Eric W. Morrison, Catalina Salgado-Salazar, Matt T. Kasson

## Abstract

*Neonectria ditissima* and *N. faginata* are canker pathogens involved in an insect-fungus disease complex of American beech (*Fagus grandifolia*) commonly known as beech bark disease (BBD). In Europe, both *N. ditissima* and *N. coccinea* are involved in BBD on European beech (*Fagus sylvatica*). Field observations across the range of BBD indicate that new infections occur primarily via ascospores. Both heterothallic (self-sterile) and homothallic (self-fertile) mating strategies have been reported for *Neonectria* fungi. As such, investigations into mating strategy are important for understanding both the disease cycle and population genetics of *Neonectria*. This is particularly important in the U.S. given that over time *N. faginata* dominates the BBD pathosystem despite high densities of non-beech hosts for *N. ditissima*. This study utilized whole-genome sequences of BBD-associated *Neonectria* spp. along with other publicly available *Neonectria* and *Corinectria* genomes and *in vitro* mating assays to characterize mating type (MAT) loci and confirm thallism for select members of *Neonectria* and *Corinectria*. MAT gene-specific primer pairs were developed to efficiently characterize the mating types of additional single ascospore strains of *N. ditissima*, *N. faginata*, and *N. coccinea* and several other related species lacking genomic data. *In vitro* mating assays were used in combination with molecular results to confirm thallism. These assays also comfirmed the sexual compatibility among *N. ditissima* strains from different plant hosts. Maximum likelihood phylogenetic analysis of both MAT1-1-1 and MAT1-2-1 sequences recovered trees with similar topology to previously published phylogenies of *Neonectria* and *Corinectria*. The results of this study indicate that all *Neonectria* and *Corinectria* tested are heterothallic based on our limited sampling and, as such, thallism cannot help explain the inevitable dominance of *N. faginata* in the BBD pathosystem.

## Introduction

Beech bark disease (BBD), a canker disease complex arising from interactions among insect and fungal causal agents, has significantly impacted the health of American (*Fagus grandifolia* Ehrh.) and European (*Fagus sylvatica* L.) beech forests throughout North America and Europe over the last century (Cale et al. 2017; Ehrlich 1934; Thomsen et al., 1949). More recently, BBD has intensified in areas where historic cold temperatures have kept the disease in check, raising concerns for the impact that global climate change may have on the expansion of this disease and other plant pathogens (Dukes et al. 2009; Kasson and Livingston 2012; McCullough and Wieferich 2001; McLaughlin and Greifenhagen 2012).

Beech bark disease requires prior infestation by a non-native scale insect *Cryptococcus fagisuga* Lind., which predisposes the host’s bark tissues to subsequent invasion by one or more closely related canker fungi: *Neonectria ditissima* ([Tul. & C. Tul.] Samuels & Rossman), *N. faginata* ([Lohman, Watson, & Ayres] Castl. & Watson), and *N. coccinea* ([Pers.] Rossman and Samuels) (Houston 1994b; Thomsen et al. 1949). *Neonectria ditissima* (formerly *N. galligena* Bres.) has been implicated in BBD both in the U.S. and in Europe while *N. faginata* (formerly *N. coccinea* var. *faginata* (Pers.:Fr.) Fr. Var. Lohman, A. M. Watson, & Ayers) appears restricted to American beech in the U.S. and *N. coccinea* to European beech in Europe (Thomsen et al. 1949; Castlebury et al. 2006). *Bionectria ochroleuca* ([Schwein.] Schroers & Samuels) has also been implicated in the U.S., but its role in BBD is not well understood (Houston et al. 1987).

*Neonectria ditissima* is perhaps best known as the causal agent of perennial target canker on many non-beech hosts including birch, maple, and walnut among others (Lohman and Watson 1943; Spaulding et al. 1936; Booth 1967). *Neonectria faginata* is unique to the BBD pathosystem in that it has only been observed causing annual cankers on American beech trees following *C. fagisuga* infestation (Castlebury et al. 2006). Other native plant hosts of *N. faginata* have not been detected. Unlike *N. ditissima* and *N. faginata*, *N. coccinea* is known to persist endophytically (asymptomatically) in the bark of European beech, with the ability to initiate disease following wounding, including but not limited to damage inflicted by *C. fagisuga* (Chapela and Boddy 1988; Hendry et al. 2002). When present, fruiting structures of *Neonectria* species are easily recognizable as bright red or orange, globose sexual ascocarps (perithecia) bearing uniseptate, hyaline ascospores. While perithecia are often products of mating between two distinct thalli of the opposite mating type, self-fertility (homothallism) can lead to the completion of the sexual cycle through selfing, which has been previously confirmed for members of the Nectriaceae (Alexopolous et al. 1996; Yun et al. 2000).

Sexual reproduction in Ascomycetes is generally understood to be regulated by the presence of one or both mating-type (MAT) idiomorphs (MAT1-1 and MAT1-2) at a mating type locus (Coppin et al. 1997; Turgeon 1998). The term “idiomorph” refers to sequences which encode different functional proteins but are found occupying the same locus in different strains. For heterothallic ascomycetes, three genes (MAT1-1-1, MAT1-1-2, and MAT1-1-3) are commonly found at the MAT locus for the MAT1-1 mating type while two genes (MAT1-2-1 and MAT1-2-2) often occur at this locus for the MAT1-2 mating type (Coppin et al. 1997; Pöggeler and Kück 2000).

These mating type idiomorphs encode polypeptides responsible for the regulation of the sexual mating cycle in filamentous fungi (Kronstad and Staben 1997). For the MAT1-1 idiomorph, MAT1-1-1, MAT1-1-2, and MAT1-1-3 encode proteins with an α domain, a PPF domain, and an HMG (high-mobility group) domain, respectively (Debuchy et al. 2010). Of these, the α domain protein encoded by MAT1-1-1 is responsible for MAT identity and sexual development (Saupe et al. 1996). The PPF and HMG domain proteins appear to have interdependent roles in fertility as the deletion of one or the other has no apparent effect, while the deletion of both has been shown to decrease fertility (Ferreira et al. 1998).

For the MAT1-2 idiomorph, MAT1-2-1 encodes a protein with a HMG domain responsible for the establishment of MAT identity (Chang and Staben 1994; Coppin et al. 1997). MAT1-2-2 encodes a small open reading frame (ORF) that does not have an apparent function (Pöggeler and Kück 2000), but this ORF appears to be absent in some filamentous fungi (Debuchy and Coppin 1992). Both MAT proteins act as transcription factors and are required for the initiation and regulation of the sexual cycle. Self-fertile (homothallic) fungi contain both MAT genes critical for sexual development (MAT1-1-1 and MAT1-2-1) at the mating type locus. In this case, reliance on a complementary mating type to complete the sexual cycle is not required.

Both homothallism and heterothallism have been previously reported for *N. ditissima* (El-Gholl et al. 1986; Krϋger 1973) while only heterothallism has been reported for *N. faginata* (Cotter and Blanchard 1978). However, these determinations relied solely on culture-based observations via *in vitro* mating assays. No molecular characterization of the MAT locus has been completed for any member of *Neonectria* despite the publicly available genomes of several species in the genus (Salgado-Salazar and Crouch 2019; Gómez-Cortecero et al. 2015; Deng et al. 2015).

Confirming the thallism of the BBD pathogens is important for several reasons: 1) thallism is predicted to affect expected patterns of genomic diversity via obligate or facultative outcrossing (Glémin and Galtier 2012); and 2) relative rates of ascospore production among species linked to potential differences in thallism could influence patterns of dominance in the BBD pathosystem and/or the perception of dominance where species-level determination are primarily made using ascospores from field-collected perithecia. In addition, MAT genes have been demonstrated as highly useful for determining phylogenetic relationships among species within a genus or clade (Lopes et al. 2018; O’Donnell et al. 2004; Turgeon 1998).

Characterizing thallism can inform our understanding of disease cycle dynamics including propagule dissemination and mode of infection. One study investigating the production and dissemination of spores by *N. ditissima* infecting yellow birch (*Betula alleghaniensis* Britton) determined ascospores to be the dominant spore type in the environment throughout the year (Lortie and Kuntz 1963). Additionally, ascospores of a related species *Corinectria fuckeliana* (C. Booth) C. González & P. Chaverri (formerly *Neonectria fuckeliana*) have also been shown to be dominant relative to asexual conidia when infecting *Pinus radiata* D. Don in New Zealand (Crane et al. 2009). Together, these results indicate the progression of diseases caused by *Neonectria* and *Corinectria* fungi may depend on ascospore production and dissemination given the proposed limited dissemination of asexual conidia by these fungi (Lortie and Kuntz 1963; Crane et al. 2009).

In addition to the potential importance of mating strategy on disease epidemiology, fungal population density assessments that depend on fruiting structure detection can be influenced by the relative rates of reproduction. Rates of visual detection are also likely influenced by fruiting structure type, where bright red perithecia are far easier to see and positively identify than the small, whitish conidiophores and sporodochia. Nearly all BBD studies that investigated interactions among these two *Neonectria* fungi have depended on the sampling and processing of perithecia to measure ascospores directly or culture the associated fungi for identification (Houston 1994; Kasson and Livingston 2009). These studies have indicated *N. faginata,* over time, supplants *N. ditissima* as the dominant pathogen in the BBD pathosystem. As with differences in optimal abiotic conditions (temperature, relative humidity) or seasonality of fruiting, differences in thallism – if they exist – could likewise influence the frequency of perithecia production and therefore, detection rates. Thus, determining thallism for *N. ditissima, N. faginata* and *N. coccinea* could enhance our ability to interpret patterns of relative abundance in the BBD system.

The objectives of this study were as follows: 1) Determine thallism among members of *Neonectria* with emphasis on BBD-associated fungi: *N. ditissima*, *N. faginata,* and *N. coccinea*. This is important as studies to determine the mating strategies of *Neonectria* fungi are limited. Furthermore, genomic data for many *Neonectria* spp. are lacking, and therefore, more general primers would be useful to permit broader characterization of mating type genes across species. 2) Confirm existing intra- and interspecies mating barriers using *in vitro* pairing assays. This is important as *N. ditissima* is reported from many plant hosts and the compatibility of strains from these various hosts remains unclear. Together, these results offer insight into mating regulation of *Neonectria* and allied fungi, thus providing an enhanced understanding of gene flow within and outside of the BBD system.

## Materials and Methods

### Genome sequencing and identification of MAT loci

Two genomes for *N. faginata* isolates (SK113 and MES1_34.1.1) were produced in association with this work and allowed for MAT gene discovery (Table 1). Full genome details will be provided in a forthcoming publication (Morrison and Garnas unpublished). Genomic data for the two *N. faginata* isolates of putatively opposite mating types as determined using the *in vitro* pairing assay described below were generated by a combination of Oxford Nanopore Technologies (ONT) and Illumina HiSeq sequencing. Genomic DNA was extracted from a *N. faginata* MAT1-1 isolate using a CTAB-chloroform DNA extraction method (van Diepen et al. 2017) and a MAT1-2 isolate using a Wizard^®^ kit (Promega, Madison, WI, USA) and suspended in 75 µl Tris-EDTA (TE) buffer (Amresco, Solon, OH, USA) with RNAse treatment to remove co-extracted RNA. The MAT1-1 isolate was sequenced using a MinION sequencer (Oxford Nanopore Technologies MIN-101B) using the unmodified 1D factory protocol (SQK-LSK109 protocol version DE_9062_v109_revD_23May2018) and a MIN-106 flow cell (FLOW MIN-106 R9 version). Both MAT1-1 and MAT1-2 isolates were subjected to Illumina HiSeq 2 x 250 PE sequencing at the University of New Hampshire Hubbard Center for Genome Studies. The ONT signal-level data was translated to FASTQ files using the Albacore v. 2.3.4 ONT proprietary basecaller resulting in >960,000 reads of which 50% were greater than 1 Kb. ONT reads were quality controlled and assembled using the Canu assembler v. 1.8 (Koren et al. 2017) with initial genome size estimate of 45 Mb. Signal level ONT data was used to polish the assembly to correct major assembly errors using Nanopolish v. 0.10.2 (Loman et al. 2015). Illumina HiSeq data was trimmed for adapter sequences and quality filtered using BBDuk (BBMap v. 38.58; Bushnell B, sourceforge.net/projects/bbmap/) resulting in 2.5 million and 11.8 million high-quality paired-end sequence reads for the MAT1-1 and MAT1-2 isolates, respectively. The MAT1-1 isolate Illumina sequences were subsequently used to further polish the Canu assembly using Pilon v. 1.22 (Walker et al. 2014). The MAT1-2 isolate sequence reads were assembled using SPAdes 3.13.1 with default settings. The resulting assemblies were assessed for contiguity using QUAST (Gurevich et al. 2013) and were checked for universal single-copy ortholog content using BUSCO v. 3.0.0 (Simão et al. 2015) with lineage Sordariomycetes. Assembly summary statistics are presented in Table S1, and complete code is available on request. Draft genome sequences are available on request.

Additionally, draft genomes of one European *N. ditissima* isolate (GenBank accession: LKCW01000000) (Gómez-Cortecero et al. 2015) and two New Zealand *N. ditissima* isolates (GenBank accessions: LDPK00000000; LDPL01000000) (Deng et al. 2015) were used to identify putative *N. ditissima* MAT idiomorphs. An unpublished *N. coccinea* draft genome (GenBank accession WPDF00000000, Castlebury et al. unpublished) was also used in this study.

To locate MAT loci within these genomes, we used NCBI GenBank tBLASTn algorithms with predicted MAT amino acid sequences derived from available MAT1-1 and MAT1-2 nucleotide sequences of two *Fusarium* (Nectriaceae) NCBI accessions: *Fusarium anguioides* (MH742713) and *Fusarium tucumaniae* (KF706656), respectively. Contigs containing sequences with an arbitrarily chosen similarity cutoff equal to or greater than 50% were selected for further examination.

### Characterizing the structure of MAT loci

Genomic data were used to create genetic maps of the MAT1-1 and MAT1-2 loci for *Neonectria ditissima* and *N. faginata* as well as the MAT1-2 locus for *N. coccinea* (Figure 1). AUGUSTUS 3.3.1 (Stanke et al. 2008) was used to predict potential gene coding regions and their resulting amino acid sequences within putative MAT loci as well as to search for conserved genes within flanking regions up to 15,000 bp upstream and downstream of the MAT idiomorphs. The selected reference genome for this prediction was *Fusarium graminearum*, which is embedded in the AUGUSTUS software. NCBI GenBank BLASTp search algorithms were used identify genes by comparing predicted amino acid sequences to the NCBI protein database.

**Figure 1:**
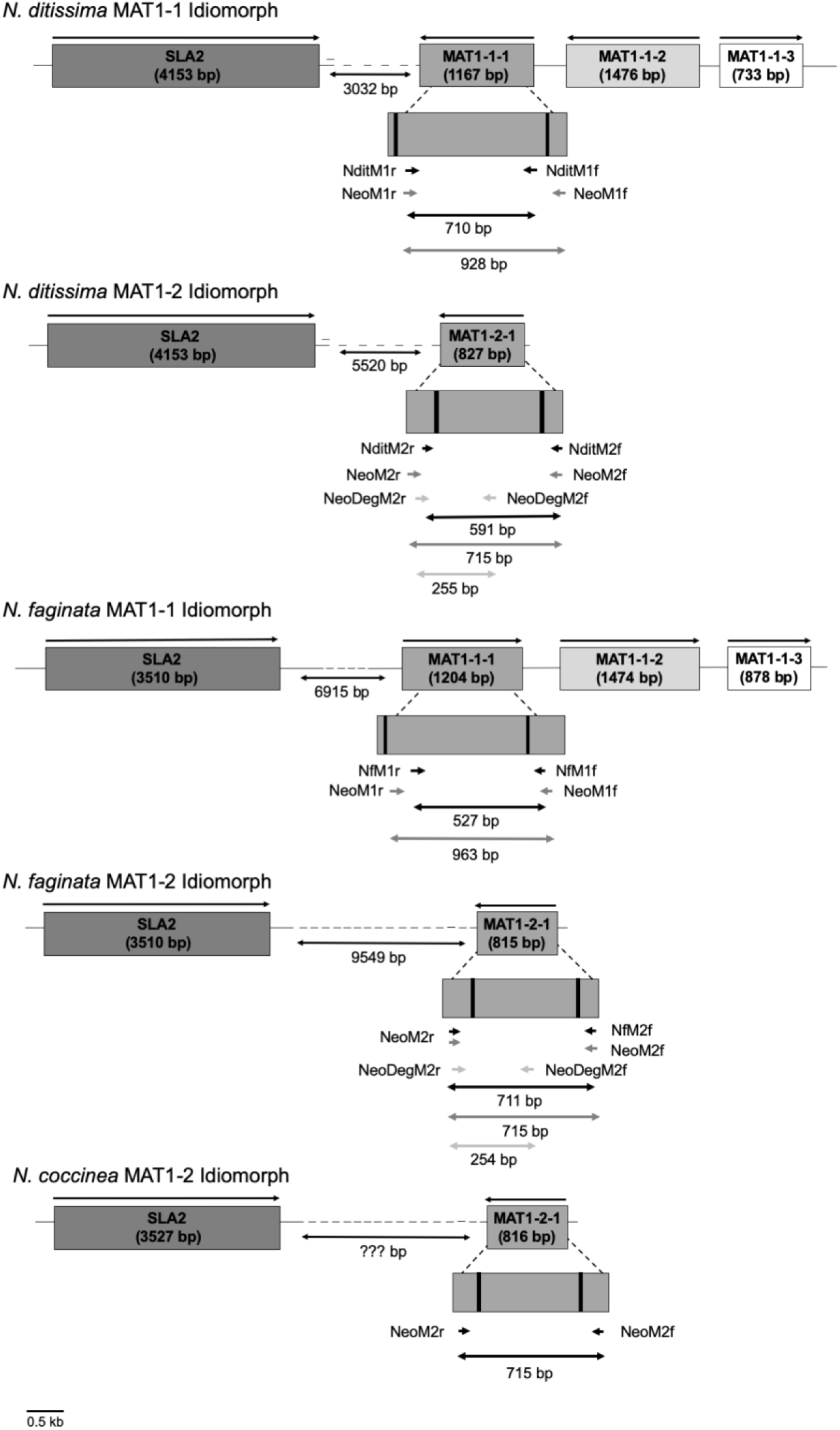
Structure of the MAT1-1 and MAT1-2 loci of the heterothallic fungi: *Neonectria ditissima*, *Neonectria faginata,* and *N. coccinea* (MAT1-2 only). Arrows above the genes indicate the 5’ – 3’ orientation. Coding sequence lengths are included below each gene identifier. For MAT1-1-1 and MAT1-2-1 genes, introns are represented by vertical black lines, and approximate primer binding locations are illustrated for each primer pair below the gene illustration. The approximate amplicon size for each primer pair is shown below the primer binding locations and is shaded based on each primer pair. All distances and sizes are estimations and not drawn to scale.

### Species-specific MAT 1-1-1 and MAT 1-2-1 primer design and PCR amplification

Once we had identified the MAT locus in each genome, we designed forward and reverse MAT1-1-1 and MAT1-2-1 specific primers for rapid characterization of mating type for *N. ditissima* and *N. faginata* isolates (Table 2). Excluding the primer binding sites, amplicon lengths were 710 bp (MAT1-1-1) and 591 bp (MAT1-2-1) for *N. ditissima.* Amplicon lengths for *N. faginata* were 527 bp (MAT1-1-1) and 612 bp (MAT1-2-1) (Table 2). Primers were manually designed in polymorphic regions distinct to each *Neonectria* species with limited repeats and approximately 60% G/C content. Primer dimer and hairpin formation among primer pairs was assessed using AutoDimer (www-s.nist.gov/dnaAnalysis; Vallone and Butler, 2004). Melting temperatures were calculated using OligoAnalyzer Tool (Intergrated DNA Technologies, Coralville, IA, USA) for standard Taq polymerase. PCR products were generated in 25 µl reactions containing 12.5 µl Bioline PCR Master Mix (Bioline USA Inc, Taunton, MA), 10.0 µl MG H2O, 1.5 µl purified DNA, and 1.0 µl of both MAT1-1 or MAT1-2 primers (25 nM; Integrated DNA Technologies, Coralville, IA, USA). PCR conditions are outlined in Table 2 for each primer set.

### Genus-level MAT 1-1-1 and MAT 1-2-1 primer design and PCR amplification

Genus-level primer pairs intended to amplify MAT1-1-1 (NeoM1f and NeoM1r) and MAT1-2-1 (NeoM2f and NeoM2r) for all included *Neonectria* species were designed as described above using available *Neonectria* genomic data (Table 2). Primer development for *Neonectria* spp. was completed using MAT1-1-1 and MAT1-2-1 sequences derived from *N. ditissima, N. faginata, N. coccinea* (MAT1-2-1), *N. punicea* ([J.C. Schmidt] Castl. & Rossman) (MAT1-1-1; GenBank accession: QGQA00000000.1), and *N. hederae* genomes ([C. Booth] Castl. & Rossman) (MAT1-2-1; GenBank accession: QGQB00000000.1). MAT1-1-1 and MAT1-2-1 nucleotide sequences were separately aligned using CLUSTAL-W (Larkin et al., 2007) within MEGA v10.1.7 (Stecher et al. 2018), and primers were designed within conserved regions to potentially increase the utility of these primers for other *Neonectria* and allied fungi. MAT1-2-1 exhibited a higher level of polymorphism among the included *Neonectria* species and therefore, a second set of degenerate primers were designed to amplifying the MAT1-2-1 gene (Table 2; NeoM2Df and NeoM2Dr). All the sequences generated were deposited in GenBank (Table 1).

### Selection of Isolates

Single ascospore-derived isolates of *N. ditissima* and *N. faginata* recovered from various geographic locations and host substrates (Table 1) were generated by squashing a single perithecium in 1ml of sterile water within a 1.5 ml Eppendorf tube, vortexing for 15 seconds to disperse ascospores, and spreading 100 µl onto a glucose-yeast extract (GYE) medium. Five germinating ascospores were transferred to a new GYE plate using a sterile needle after 24 hours and were allowed to grow for one week before replating. Each isolate was grown on GYE for two weeks and then identified using colony morphology based on type descriptions (Castlebury et al., 2006).

Genomic DNA was extracted from isolates or directly from perithecia as described above. For each species identified morphologically, three random isolate identifications were confirmed by sequencing the ribosomal internal transcribed spacer region (ITS) using primers ITS5 (5’ – GGAAGTAAAAGTCGTAACAAGG – 3’) and ITS4 (5’ – TCCTCCGCTTATTGATATGC – 3’) (White et al., 1990). The PCR protocol was as follows: 95 °C for 2 min followed by 35 cycles of 95 °C for 30 sec, 56 °C for 30 sec, 72 °C for 1 min with a final extension at 72 °C for 7 min. EF1-α sequencing was completed using primers EF1728F (5’ – CATCGAGAAGTTCGAGAAGG – 3’) (Carbone and Kohn, 1999) and EF1-1567R (5’ – ACHGTRCCRATACCACCRATCTT – 3’) (Rehner 2001) with the following PCR protocol: 95 °C for 2 min followed by 35 cycles of 95 °C for 30 sec, 55 °C for 1 min, 72 °C for 1 min with a final extension at 72 °C for 7 min.

PCR products were visualized with gel electrophoresis by adding 4 μl SYBR gold (Invitrogen, Grand Island, NY, USA) and 4 μl loading dye (5Prime, Gaithersburg, MD) to products. Samples were then loaded into a 1.5%, wt/vol, agarose gel (Amresco, Solon, OH, USA) made with 0.5% Tris-borate-EDTA buffer (Amresco, Solon, OH, USA). To compare sizes, 100-bp and 1-kb molecular ladders (Omega Bio-tek, Norcross, GA, USA) were also added to gels. Electrophoresis was performed at 90V for 75 minutes. Bands were visualized on a UV transilluminator (Syngene, Frederick, MD, USA).

Positive reactions were purified using ExoSAP-IT (Affymetrix, Santa Clara, CA, USA) according to the manufacturer’s recommendations. Purified PCR products were Sanger sequenced in forward and reverse directions using the same PCR primers (Eurofins, Huntsville, AL, USA). BLASTn searches were then used to identify species based on the best match in the NCBI database.

### Mating type gene screening

All selected *N. ditissima* and *N. faginata* (22 and 18 isolates, respectively) were screened for the presence of MAT1-1-1 and/or MAT1-2-1 using species-specific and genus-level primer sets with the PCR protocols listed in Table 2. All PCR products were visualized, and a subset of positive reactions were sequenced as described for ITS and EF1-α amplicons.

To confirm the specificity of species-specific primer sets, mating type PCR reactions were performed using *N. ditissima* MAT primers for *N. faginata* isolates and vice versa. Additionally, a number of other members of the Nectriaceae were also tested using both *N. ditissima* and *N. faginata* MAT primers as described above. These included isolates of *N. coccinea, N. neomacrospora* ([C. Booth & Samuels] Mantiri & Samuels)*, Fusarium concolor* (Reinking, 1934)*, Nectria magnoliae* (M.L. Lohman & Hepting, 1943) and an additional unresolved species for which additional data is needed to confirm identity: *Corinectria* aff. *fuckeliana* (99.37% EF1-α sequence similarity to NCBI Genbank accession MK911707.1) (Table 1). Genus-level *Neonectria* MAT primer sets were similarly tested using representatives from all five Nectriaceae species tested in this study.

All resulting sequences were aligned as described above and compared to genome-derived MAT1-1-1 or MAT1-2-1 sequences to confirm their identity based on a sequence similarity. All sequences having greater than 70% sequence similarity were selected for further analysis. This threshold was determined by comparing MAT1-1-1 and MAT1-2-1 genome-derived sequences from *N. ditissima* to the more distantly related *Fusarium anguioides* (MAT1-1-1; Genbank Accession: MH742713) and *Fusarium tucumaniae* (MAT1-2-1; Genbank Accession: KF706656) sequences.

### Phylogenetic analysis and protein alignment

To examine evolutionary patterns and divergence in mating type genes, we constructed phylogenetic trees using the mating type gene sequence data produced in this study together with comparable sequences for other Nectriaceae available in NCBI Genbank, including 11 strains representing 7 species (Table 1). All analyses were completed using MEGA v10.1.7 (Stecher et al. 2018). MAT nucleotide sequences were aligned using CLUSTAL-W (Larkin et al. 2007) and the best-fit nucleotide substitution model was chosen using Model Test AICc scores in MEGA v10.1.7. MAT1-1-1 and MAT1-2-1 maximum-likelihood trees were constructed independently using the Kimura 2-parameter model with gamma distribution (K2+G) and 1000 bootstrap replicates. For both trees, MAT1-1-1 and MAT1-2-1 sequences from *Ophiocordyceps xeufengensis* and *Ustilaginoidea virens* served as outgroup taxa (Table 1).

We performed protein alignments to characterized divergence among species that could play a role in the maintenance of mating barriers. Protein sequences were predicted from one MAT1-1-1 and one MAT1-2-1 coding sequence from each species using ExPASy-Translate tool (https://web.expasy.org/translate/). Resulting protein sequences were aligned using CLUSTAL-W within MEGA v10.1.7. Boxshade Server v.3.21 (https://embnet.vital-it.ch/software/BOX_form.html) was used to visualize shared amino acids within each sequence.

### In vitro mating assay

To demonstrate mating among MAT1-1 and MAT1-2 strains, an *in vitro* mating assay was performed with six MAT1-1-1 and six MAT1-2-1 isolates of both *N. faginata* and *N. ditissima*. These isolates varied in geographic origin and for *N. ditissima*, host substrate as to test compatibility among representatives from allopatric populations and potentially host-specific *N. ditissima* isolates. This pairing assay did not include *N. coccinea* as only genomic DNA was available. Selected isolates were grown on glucose-yeast extract agar for two weeks. Each selected isolate was then paired three times with itself, with an isolate of the same mating type, and with two isolates of the opposite mating type for a total of twelve pairings per isolate. For each pairing, a 5x5-mm fungus-colonized agar plug from each isolate was placed on opposite sides of a sterile, flat toothpick placed atop the media along the center point of the petri dish (Figure 2A). All plates were parafilmed and placed at 20 °C with a 16 h/8 h light/dark cycle under cool fluorescent lamps. Plates were checked weekly for perithecia formation for up to 12 weeks (Figure 2B).

**Figure 2:**
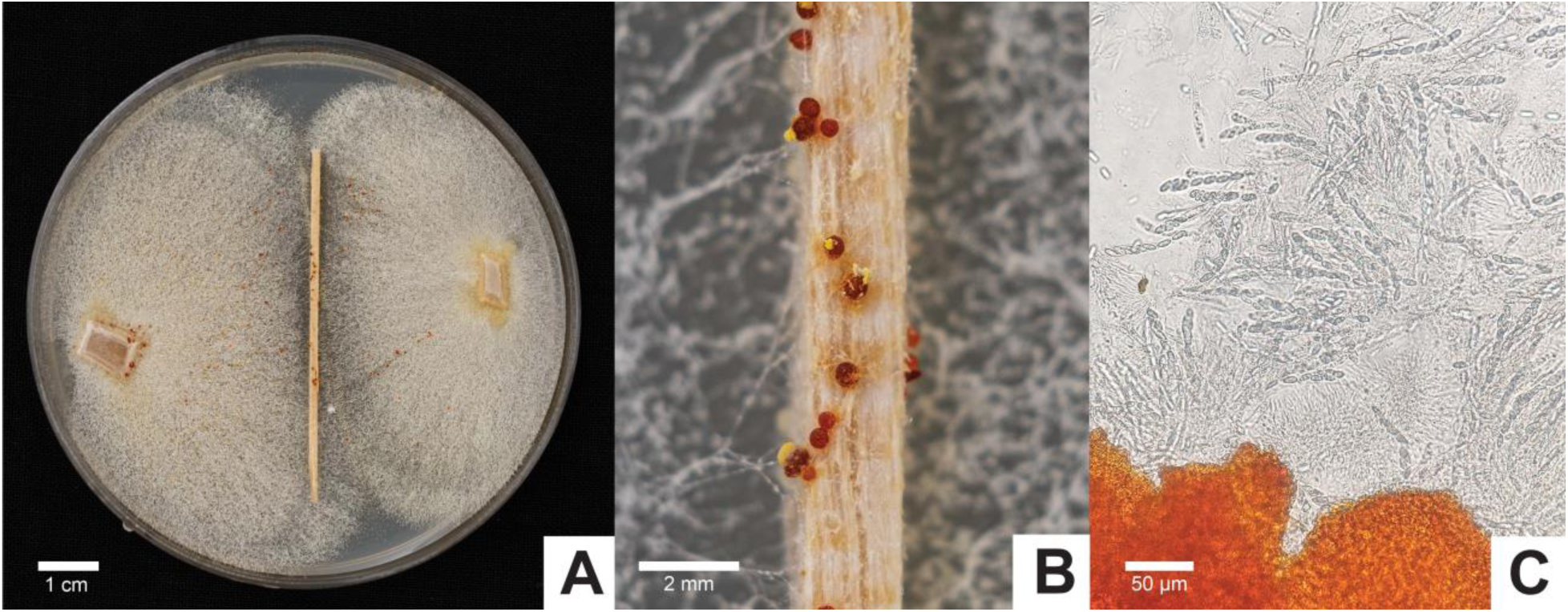
*In vitro* mating assay for *Neonectria* spp. and related fungi: A) Approximately 5mm x 5mm colonized plugs of confirmed MAT1-1 and/or MAT1-2 strains placed on either side of a sterile wooden toothpick placed upon GYE media; B) Perithecia exuding ascospores on toothpick after approximately 12 weeks; C) Perithecia squash mount showing ascospores within asci. *Nectria magnoliae* shown in figure. Individual scale bars are included in each panel.

As perithecia were produced, squash mounts were used to check for ascospores using light microscopy (Figure 2C). Additionally, ascospores were checked for viability by removing a single perithecium, macerating in 1 ml of sterile dH2O, spreading 300 µl of the spore suspension on GYE, and observing growth. For one progeny plate from each paired isolate set, ten germinating ascospores were sub-cultured onto a new GYE plate after 24 hours and allowed to grow for one week. Genomic DNA was extracted from the ten subcultured progeny and screened for MAT genes as described above. Given that progeny should segregate 1:1 for MAT1-1 and MAT1-2, all ascospore suspension plates were incubated for up to 12 weeks to observe mating among the progeny.

A subsequent interspecies mating assay was performed to test mating capability among *N. ditissima, N. faginata,* and other Nectriaceous fungi. Here, MAT1-1- and MAT1-2-positive isolates of *Neonectria ditissima* (MAT1-1: NdBl001; MAT1-2: NdSam003) and *N. faginata* (MAT1-1: NfFg005; MAT1-2: NfFg008) were paired three times each with an isolate of *Nectria magnoliae* (isolate no. NecmLt005) and *Corinectria* aff. *fuckeliana* (isolate no. CafPr004) as described above. Additionally, both *Neonectria ditissima* isolates were paired three times with the *N. faginata* isolate of the opposite mating type. Plates were weekly checked for perithecia formation for up to 12 weeks and processed as described above.

## Results

### Identification and structure of MAT loci in N. ditissima and N. faginata

Each of the *N. faginata* and *N. ditissima* genomes contained either the MAT1-1 or MAT 1-2 idiomorph, and the single *N. coccinea* genome contained only a MAT1-2 idiomorph. MAT genes of the opposite mating type were not found within the genomes. AUGUSTUS analyses of the MAT loci and flanking genes revealed a similar genetic structure for *N. coccinea*, *N. ditissima* and *N. faginata* with only minor differences in the MAT gene open reading frame (ORF) and intron lengths (Figure 1). The MAT1-1-1 ORF for *N. ditissima* was 1,167 bp (357 amino acids [aa]; GenBank accession XXXXX) with two introns of 48 bp and 46 bp, while the MAT1-1-1 ORF for *N. faginata* was 1204 bp (371 aa; GenBank accession XXXXX) with two introns of 46 bp and 44 bp. The MAT1-2-1 ORF for *N. ditissima* was 827 bp (243 aa; GenBank accession XXXXX) with two introns of 48 bp and 49 bp, while the ORF for *N. faginata* was 815 bp (239 aa; GenBank accession XXXXX) with two introns of 47 bp and 50 bp. Additionally, the genetic structure of MAT1-2-1 of *N. coccinea* was found to be similar to *N. ditissima* and *N. faginata* with an ORF of 816 (238 aa; GenBank accession XXXXX) with two introns of 50 bp and 48 bp.

Two commonly co-occurring MAT associated genes were found near MAT1-1-1 in both *N. ditissima* and *N. faginata,* including MAT1-1-2 and MAT1-1-3 (Figure 1; Coppin et al. 1997). For each of the three *Neonectria* spp., evidence of co-occurrence of MAT1-2-1 and MAT1-2-2 was not found. Both *N. ditissima* and *N. faginata* MAT loci were flanked by the conserved SLA2 gene previously described as being associated with MAT loci (Debuchy and Turgeon 2006). SLA2 was found to occur in a separate contig of the *N. coccinea* de novo genome assembly, and although present, could not be shown to be part of the MAT locus without further assembly. APN2 is a second conserved gene often flanking the MAT loci of other Ascomycetes (Debuchy and Turgeon 2006), but this gene was not identified in the regions flanking the MAT loci using the de novo genome assemblies analyzed in this study.

### Species specific MAT1-1-1 and MAT1-2-1 primers

*Neonectria ditissima* (NdM1f/r) and *N. faginata* (NdM2f/r) MAT primer pairs amplified a single product of the expected size for their target species (Table 2). Sequencing confirmed the identity of all PCR products which exhibited 99% or greater sequence identity with the target MAT sequences. All of the 160 (40 initially screened or 120 progenies from *in vitro* crosses) single-spore derived isolates tested yielded either MAT1-1-1 or MAT1-2-1 amplicons while DNA extractions from perithecia (N = 7) containing ascospores of both mating types yielded both MAT products. Species-specific primers designed for *N. faginata* did not amplify either MAT gene in *N. ditissima.* Likewise, *N. ditissima-*specific primers did not amplify either MAT gene in *N. faginata*.

MAT1-1-1 and MAT1-2-1 primer pairs designed for *N. ditissima* did not amplify DNA in any of the other tested species (Table 3). In contrast, the *N. faginata* MAT1-1-1 primer pair amplified MAT1-1-1 in MAT1-1 isolates of *N. coccinea, N. neomacrospora*, and *Nectria magnoliae*. In contrast, MAT1-2-1 primers for *N. faginata* did not amplify MAT1-2-1 for any other tested species.

Amplification of non-mating type associated proteins was observed when applying *N. faginata* MAT1-1-1 primers to MAT1-2 strains of *N. coccinea, Corinectria* aff. *fuckeliana*, and *N. neomacrospora*. Sequencing of these amplicons confirmed a putative MFS-type transporter protein (∼120 bp > target) in *N. coccinea* and two undescribed hypothetical proteins in *N. neomacrospora* (∼470 bp > target) and *C.* aff. *fuckeliana* (∼470 bp > target) (Table S2). Additionally, amplification of non-mating type associated proteins was observed when *N. faginata* MAT1-2-1 primers were applied to MAT1-1 *N. coccinea* and *C.* aff. *fuckeliana* isolates including hypothetical proteins in *N. coccinea* (∼540 bp > target) and *C.* aff. *fuckeliana* (∼190 bp > target) (Table S2).

### Genus-level Neonectria MAT1-1-1 and MAT1-2-1 primers

Genus-level MAT primer pairs (NeoM1f/r and NeoM2f/r) successfully amplified both MAT1-1-1 and MAT1-2-1 gene for *N. ditissima* and *N. faginata* (Supplemental Figure 1). These same primers also amplified MAT1-1-1 for *Corinectria* aff. *fuckeliana* (Table 3) but failed to amplify MAT1-1-1 and MAT1-2-1 for all other non-target fungi.

The MAT1-2-1 degenerate primer pair (NeoM1df/r) successfully amplified MAT1-2-1 for all species tested, including a more distantly related *Fusarium concolor* isolate. Non-target amplification of a hypothetical protein (∼590 bp > target) was observed when the NeoM2f/r primer pair was applied to MAT1-1 *N. coccinea* and both MAT1-1 and MAT1-2 *N. neomacrospora* isolates.

### Phylogenetic analyses and protein alignments

Phylogenetic analyses of MAT1-1-1 and MAT1-2-1 sequences resulted in similar tree topologies (Figures 3 and 4). Analysis of MAT1-1-1 grouped all *Neonectria* species into a strongly supported monophyletic clade sister to *Corinectria* (formerly *Neonectria*) (González and Chaverri 2017) (Figure 3). Within *Neonectria*, a clade containing *N. coccinea, N. faginata* and *N. punicea* was resolved as sister to *Nectria magnoliae* from *Liriodendron tulipifera* and *Magnolia fraseri*. For MAT1-2-1, all included *Neonectria* species resolved to a monophyletic clade that was sister to a clade containing *Corinectria* aff. *fuckeliana* and the three species of *Fusarium* (Figure 4). Additionally, two sister clades within *Neonectria* were resolved: 1) *N. faginata* was sister to a clade containing *N. coccinea* and *Nectria magnoliae*, and 2) *Neonectria ditissima* and *N. neomacrospora* formed a monophyletic clade sister to *N. hederae*. All isolates of *N. ditissima* formed a single lineage regardless of plant host for both MAT1-1-1 and MAT1-2-1.

**Figure 3:**
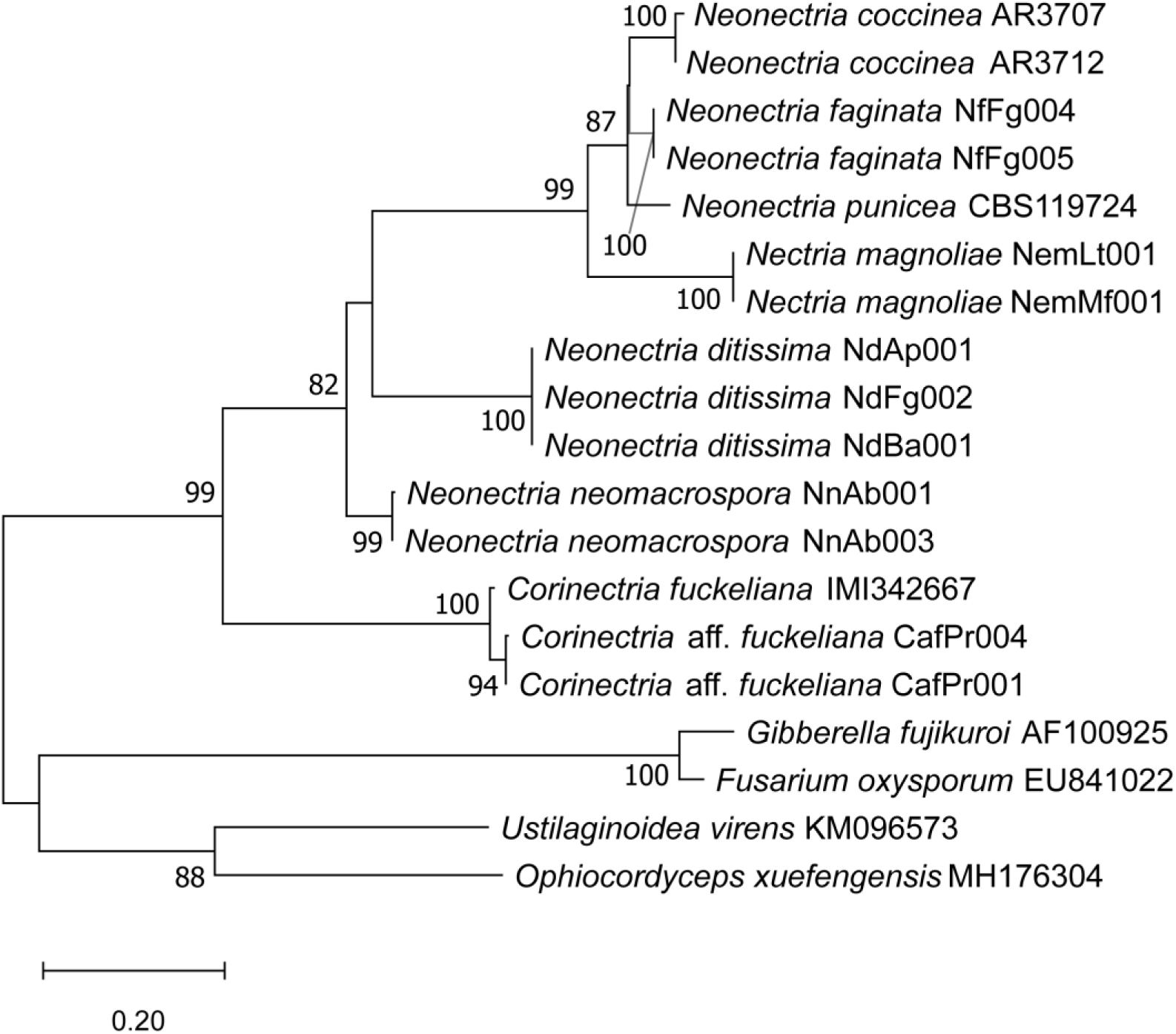
Phylogenetic relationships of *Neonectria* spp. and related fungi based on MAT1-1-1 gene sequence data. The phylogeny was inferred using a Maximum Likelihood analysis based on the Kimura 2-parameter model with gamma distribution (K2+G) and 1000 bootstrap replicates. Bootstrap values > 70% are given at the nodes. Branch lengths represent the number of substitutions per site. Outgroup includes *Ophiocordyceps xeufengensis* and *Ustilaginoidea virens*.

**Figure 4:**
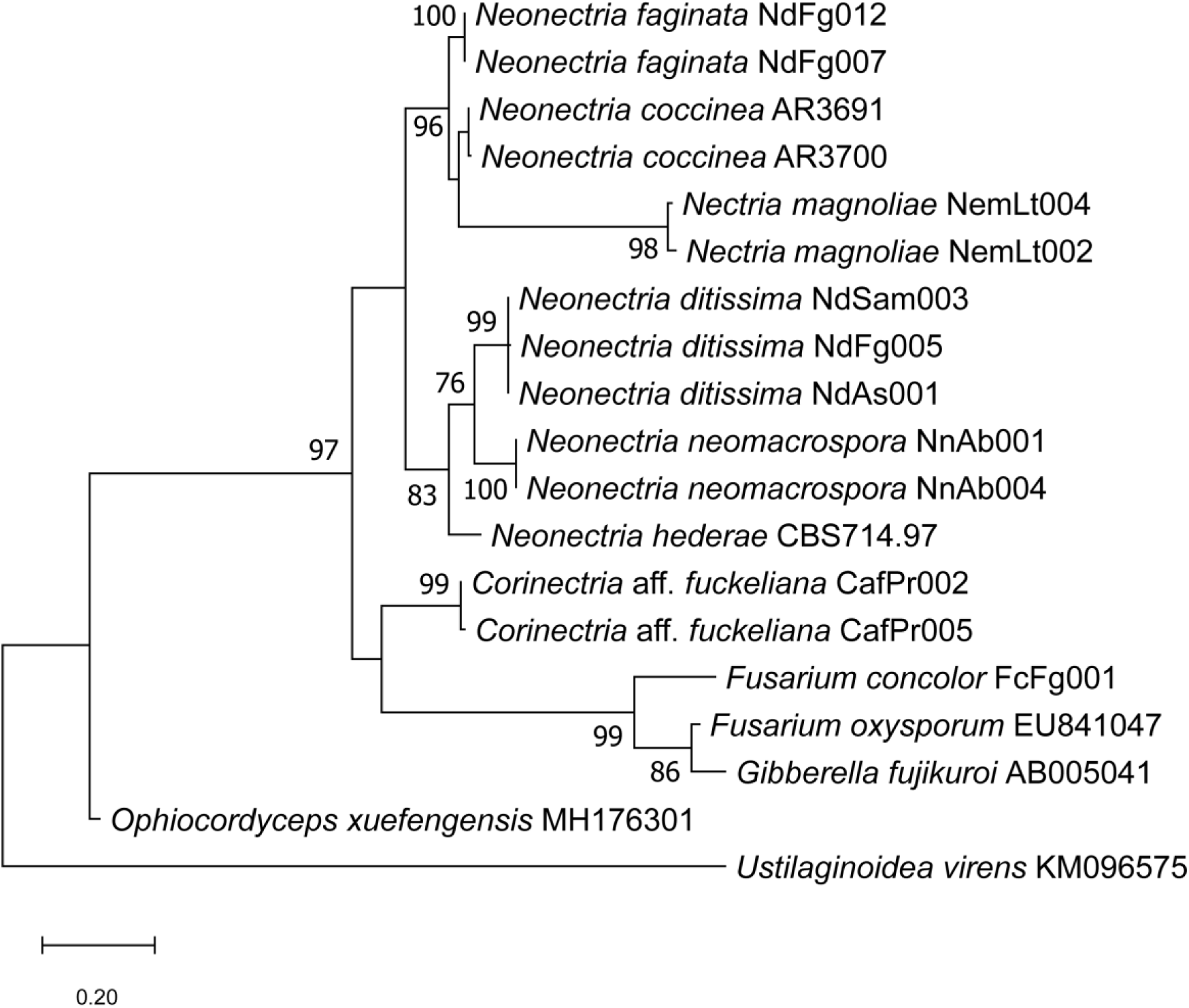
Phylogenetic relationships of *Neonectria* spp. and related fungi based on MAT1-2-1 gene sequence data. The phylogeny was inferred using a Maximum Likelihood analysis based on the Kimura 2-parameter model with gamma distribution (K2+G) and 1000 bootstrap replicates. Bootstrap values > 70% are given at the nodes. Branch lengths represent the number of substitutions per site. Outgroup includes *Ophiocordyceps xeufengensis* and *Ustilaginoidea virens*.

Protein alignments included approximately 100 amino acids for MAT1-1-1 sequences and approximately 75 amino acids for MAT1-2-1 sequences within the conserved region of these genes (Figure 5). *Neonectria coccinea, Nectria magnoliae, Corinectria* aff. *fuckeliana*, and *Fusarium concolor* were missing the first five MAT1-2-1 amino acids, and therefore, results reported for MAT1-2-1 below represent values with (full) and without (partial) the first five amino acids. Protein alignments for both MAT1-1-1 and MAT1-2-1 were most similar among *Neonectria* spp. with 28% and 29.3% (or 36% for full sequence comparisons) of amino acids conserved across all species considered, respectively. Of these shared amino acids, 19% MAT1-1-1 and 25.3% MAT1-2-1 amino acids (partial seqs) were shared with included *Corinectria* spp. Across all included fungi, only 5% of amino acids were shared among MAT1-1-1 sequences and 21.3% (or 28% for full sequence comparisons) amino acids were shared among MAT1-2-1 sequences.

**Figure 5:**
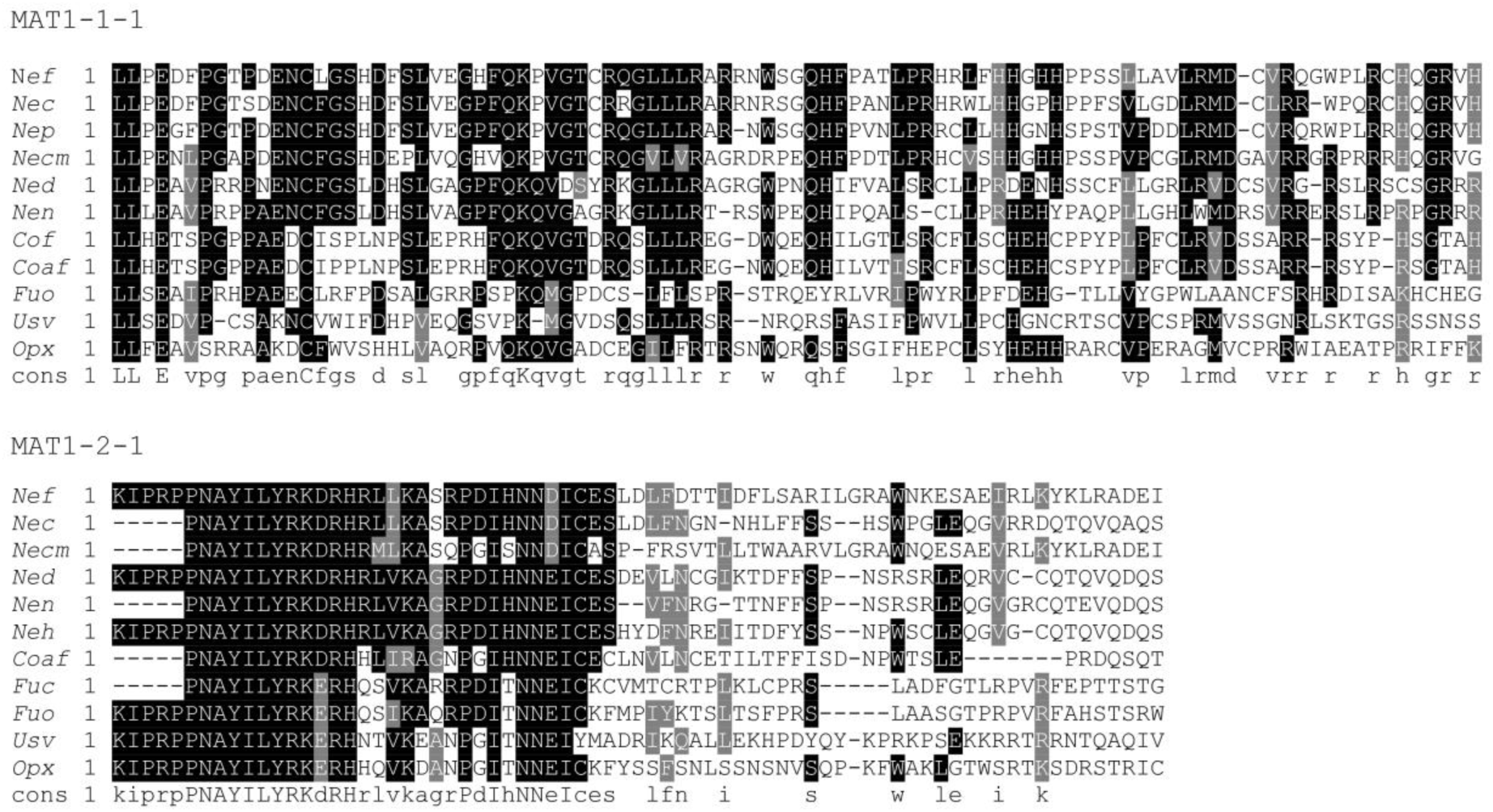
MAT1-1-1 and MAT1-2-1 amino acid alignments including a selection of study fungal species. Name abbreviations are as follows: *Nef = Neonectria faginata*; *Nec = Neonectria coccinea*; *Nep = Neonectria punciea*; *Necm = Nectria magnoliae*; *Ned = Neonectria ditissima*; *Nen = Neonectria neomacrospora; Neh = Neonectria hederae*; *Cof* = *Corinectria fuckeliana*; *Coaf = Corinectria* aff. *fuckeliana*; *Fuo = Fusarium oxysporum*; *Usv = Ustilaginoidea virens*; *Opx = Ophiocordyceps xuefengensis;* Cons = consensus. Uppercase letters within the consensus sequence represent positions with identical amino acids. Lower case letters represent positions with similar amino acids. Black shading represents conserved amino acids among 50% or more species. Grey shading represents shared amino acids with shared characteristics/properties among species.

### In vitro pairing assay results

For *N. faginata* and *N. ditissima* intraspecies pairings, perithecia with viable ascospores were produced for all pairings between MAT1-1 and MAT1-2 strains with one exception: one pairing between isolates NfFg006 and NfFg012 failed to produce perithecia within the 12-week assay (Table 4). All ascospore suspension plates containing ascospore progeny from single, squashed perithecia resulting from MAT1-1 x MAT1-2 pairings also yielded perithecia. No pairings with an isolate of the same mating type, including self-pairings, yielded perithecia. Additionally, no evidence of mating was observed in interspecies pairings.

Each progeny included in the subsequent PCR screening resulted in the amplification of a single MAT gene. Both MAT genes were found to be segregated among progeny at a ratio not significantly divergent from the expected 1:1 for *N. ditissima* (χ*^2^* = 0.07, df = 1,60; *P* = 0.80) and *N. faginata* (χ*^2^* = 0.27, df = 1,60; *P* = 0.606) progeny (Table 4).

## Discussion

Beech bark disease was first reported in Halifax, Nova Scotia, Canada around 1890 and continues to spread throughout the range of American beech (Hewitt 1914; Erhlich 1934). Likewise, BBD continues to impact European Beech throughout Europe (Cicák et al. 2006; Cicák and Mihál 2008). Despite the ecological importance of this disease (Houston 1994b; Garnas et al. 2011a,b), certain aspects of BBD biology and epidemiology remain largely unexplored, including the mating strategies of *N. ditissima, N. faginata,* and *N. coccinea*. Previous studies regarding the thallism of *N. ditissima* and *N. faginata* (El-Gholl et al. 1986; Krϋger 1973; Cotter and Blanchard 1978) were either inconclusive or sample limited, and as such, the role of mating strategy in the BBD pathosystem in the U.S. has historically been uncertain. Thallism is particularly important to the BBD pathosystem given that at least two dominant, sexually reproducing pathogens are causal agents of the disease in both the U.S. and Europe. Both can colonize single trees and, in some instances, have been found to co-occur within the same 2.5” diameter bark disk (Kasson and Livingston 2009).

*Neonectria faginata* has been previously reported as the dominant pathogen in the BBD system in North America (Houston 1994; Kasson and Livingston 2009). Mating strategy may have provided one explanation for its dominance, but as shown in this study, both *N. faginata* and *N. ditissima* are heterothallic fungi based on limited sampling. Therefore, the possibility of any advantage that homothallism might confer is eliminated, indicating that *N. faginata* is likely dominant due to an increased level of virulence, other advantageous traits, or some combination of these factors.

Despite having resolved the mating strategy of these fungi, the potential overrepresentation of either *N. ditissima* or *N. faginata* in previous studies may be due to perithecia-dependent sampling bias. For example, environmental conditions (*e.g.* RH, temperature) and time required for perithecia production may significantly differ between these two fungi. Given potential seasonal differences in fruiting, sampling at a single time point in the year could potentially yield a community not representative of the relative abundance of the *Neonectria* spp. present. Additionally, casual observations of *N. ditissima* on non-beech hosts can reveal very limited to no perithecia occurring at a given infection site. This may be in part due to host susceptibility, resulting in more or less necrotic tissue, which is generally considered favorable for perithecia production by necrotrophic fungi.

Host specificity may develop through the evolution of mating barriers among strains of a single species occurring on differing hosts. Results from this study have demonstrated a lack of reproductive barriers among *N. ditissima* strains from several plant hosts. All pairings of either MAT1-1-1 or MAT1-2-1 strains from different plant hosts resulted in perithecia formation. While a lack of host specificity by *N. ditissima* has been previously demonstrated using pathogenicity assays (Lortie 1969; Ng and Roberts 1974; Barnard et al. 1988; Plante and Bernier 1997), mating among *N. ditissima* strains infecting co-occurring tree species had not been previously tested until now. Given the lack of reproductive barriers, evolution of host specificity is limited by obligate outcrossing among these co-occurring strains.

The phylogenetic analyses using MAT gene sequences were found to be in agreement with previously resolved relationships demonstrated using EF1-α, RPB2, and β-tubulin (Castlebury et al. 2006). This finding confirms the utility of MAT genes in resolving relationships among *Neonectria* and allied fungi. Additionally, *N. ditissima* isolates from different plant hosts included in the phylogeny form a single lineage providing additional evidence for the lack of reproductive barriers, limiting the possibility the development of host specificity within *N. ditissima* populations co-occurring across multiple hosts. The amino acid alignments visualized protein sequence divergence potentially contributing to mating barriers among these closely related species that were confirmed by inter-species mating assays. Using such methods to test intraspecies MAT gene diversity among these and closely allied fungi may prove valuable for broader surveys.

Limited amplification of non-mating type proteins was observed among several MAT primer – species combinations for isolates of the opposite mating type. However, in all cases, the sequences were found to be larger than the expected product size. Given that such issues can lead to erroneous conclusions, electrophoresis gels should be run to 100 bp resolution and additionally, PCR conditions should be optimized for the target species.

## Conclusion

In this study, we confirm heterothallism and characterized the MAT idiomorphs of *N. ditissima*, *N. faginata*, *N. coccinea* and several other members of Nectriaceae. These findings provide additional insight into characteristics that may shape the community and population dynamics of the beech bark disease complex and its causal agents. Additional studies are needed to further understand the fungal dynamics of *N. ditissima, N. faginata,* and *N. coccinea* in their respective BBD systems. These efforts include: 1) identifying differing environmental factors required for perithecia production among *Neonectria* spp.; 2) characterizing variability in sexual reproduction by *N. ditissima* across host substrates; 3) assessing the potential for interspecific hybridization between closely related *Neonectria* spp. found co-occurring on beech and other hosts but excluded from this study; and 4) comprehensive screening of additional isolates from populations not sampled in this study to assess intraspecies MAT gene diversity and uncover possible intraspecies mating barriers.

## Supporting information

Main and Supplemental Tables

## Acknowledgements

The authors thank Nicole Utano, Braley Burke and Amy Metheny for their assistance with field collections and sample processing. C.M.S was supported, in part, by the WVU Outstanding Merit Fellowships for Continuing Doctoral Students. M.T.K. by funds from West Virginia Agricultural and Forestry Experiment Station. Work by J.R.G. and E.W.M was supported by the USDA National Institute of Food and Agriculture McIntire-Stennis Project (Accession No. 1012453). Partial funding was provided by the New Hampshire Agricultural Experiment Station. This is Scientific Contribution Number . The mention of firm names or trade products does not imply that they are endorsed or recommended by the US Department of Agriculture over other firms or similar products not mentioned. The USDA is an equal opportunity provider and employer.

---

**Supplemental Figure 1:**
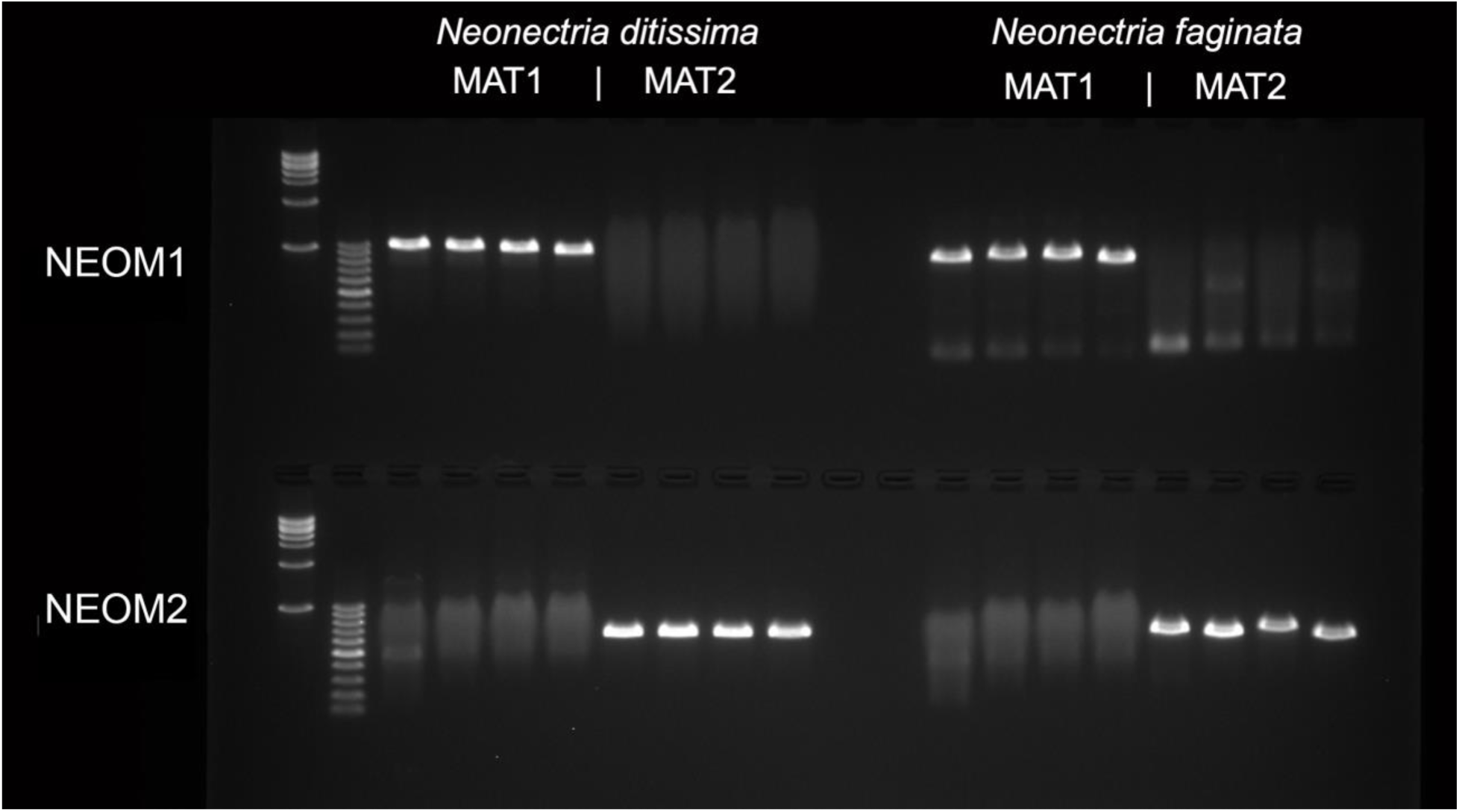
Electrophoresis gel photo demonstrating specific amplification of MAT1-1-1 and MAT1-2-1 by the genus-level primer pair (NeoM1f/r and NeoM2f/r) for both *N. ditissima* and *N. faginata*.

